# Diel dynamics of dissolved organic matter and heterotrophic prokaryotes reveal enhanced growth at the ocean’s mesopelagic fish layer during daytime

**DOI:** 10.1101/712570

**Authors:** Xosé Anxelu G. Morán, Francisca C. García, Anders Røstad, Luis Silva, Najwa Al-Otaibi, Xabier Irigoien, Maria L. Calleja

## Abstract

Contrary to epipelagic waters, where biogeochemical processes closely follow the light and dark periods, little is known about diel cycles in the ocean’s mesopelagic realm. Here, we monitored the dynamics of dissolved organic matter (DOM) and planktonic heterotrophic prokaryotes every 2 h for one day at 0 and 550 m (a depth occupied by vertically migrating fish during light hours) in oligotrophic waters of the central Red Sea. We additionally performed predator-free seawater incubations of samples collected from the same site both at midnight and at noon. Comparable in situ variability in microbial biomass and dissolved organic carbon concentration suggests a diel supply of fresh DOM in both layers. The presence of fish in the mesopelagic zone during daytime promoted a sustained, longer growth of larger prokaryotic cells. The specific growth rates were consistently higher in the noon experiments from both depths (surface: 0.34 vs. 0.18 d^−1^, mesopelagic: 0.16 vs. 0.09 d^−1^). Heterotrophic bacteria and archaea in the mesopelagic fish layer were also more efficient at converting DOM into new biomass. These results suggest that the ocean’s twilight zone receives a consistent diurnal supply of labile DOM from diel vertical migrating fishes, enabling an unexpectedly active community of heterotrophic prokaryotes.

---

Planktonic heterotrophic prokaryotes (HP) pertaining to the domains Bacteria and Archaea rely on labile dissolved organic matter (DOM) for metabolism and growth ^1, 2, 3^. In surface waters, diel cycles in HP biomass and activity have been related to the photosynthetic activity of phytoplankton ^4^, which obviously follows sunlight. Heterotrophic prokaryotes dependence on DOM derived from planktonic algae ^5^ was reported to increase offshore, far from coastal inputs, in temperate and polar ecosystems ^6^. Although this relationship, also known as bacterioplankton-phytoplankton coupling, has been the subject of debate ^7,8^, in regions with low DOM advection (e.g. at permanently stratified sites without anthropogenic or riverine inputs nearby, such as oligotrophic tropical waters), we might expect strong diel signals in the response of heterotrophic prokaryotes coupled with the activity of primary producers ^9^. In this regard, the Red Sea offers a unique opportunity to study biogeochemical processes in oligotrophic ecosystems. With no permanent rivers, the only allochthonous inputs of DOM come from urban centers such as Suez, Ghardaqa, Jeddah or Port Sudan, coastal macrophytes ^10, 11^ or dust events ^12, 13^.

While epipelagic processes driven by primary production are well known ^14, 15^, large gaps in our understanding of the ecology and biogeochemistry of the mesopelagic zone (i.e. waters between 200 m and 1000 m) remain ^16^. In the mesopelagic realm, trophic interactions between microbes and metazoa have been long neglected. The available studies have focused mostly on mesozooplankton^17, 18, 19^. However, recent reports on the large biomass contributed to the ocean’s biota by mesopelagic fishes performing diel vertical migration (DVM) suggest they may also play an important role as rapid vectors of labile organic mater^20, 21^. DVM can affect only a fraction of individuals from a given population ^20^. In the Red Sea, virtually the entire populations of mesopelagic fish migrate daily between the surface and the so-called deep scattering layer (DSL) located at 400-650 m in the mesopelagic zone ^22, 23^. DVM fishes have been recently suggested to generate hotspots for heterotrophic prokaryotes, yielding significantly higher bacterial growth efficiencies compared with shallower layers ^24^. An analysis of a 24 h intensive sampling at the same location has supported the existence of diel inputs of labile DOM fueling the HP community at the depths occupied by mesopelagic fishes during daytime ^25^. Both DOC concentrations and high nucleic acid content (HNA) bacteria and archaea, usually made up of copiotrophic taxa ^26, 27^ and more active than the low nucleic acid content (LNA) group^28, 29, 30^, fluctuated as widely in waters below 200 m as in the upper layers. However, for the hypothesis of the mesopelagic labile DOM hotspots to be true, we should be able to demonstrate that the presence or absence of fishes in the twilight zone does make a difference.

Here, we report on the results of two short-term incubations with water collected from the epi- and mesopelagic layers (surface and 550 m, respectively) of the central Red Sea at midnight and at the following midday. After removing protistan grazers and other larger organisms by filtration, we followed the dynamics of DOM-heterotrophic prokaryotes interactions for 8 days. In parallel, we conducted a high frequency (every 2 h for a full 24 h starting at noon) characterization of the same depths focusing on the response of heterotrophic prokaryotes abundance, cell size and biomass to changes in DOM concentrations including its fluorescent properties, previously unreported for this basin. The specific objectives of this study were: i) to assess the diurnal scales of variability in the standing stocks of HP and DOM in epipelagic and mesopelagic waters of the central Red Sea, and ii) to test for differences in the specific growth rate, maximum biomass and growth efficiency of HP between nighttime and daytime in both layers. Our hypothesis is that DOM supplied by DVM fishes in the mesopelagic zone during the day had a commesurable effect on the above-mentioned variables.

## Results

### Environmental variability of DOC and heterotrophic prokaryotes

The complete diel vertical migration of the mesopelagic fishes present at the study site can be clearly seen in the echogram of **Fig. 1**, with the deeper, more intense layer (dominated by Benthosema pterotum) occupying the depths between ca. 520 and 630 m during daytime on March 6^th^ 2016. **Fig. 2** shows the diel variability of DOC concentrations and the biomass of HP at the station’s surface and 550 m depth. Mean DOC values were almost 50% higher at 0 than at 550 m (71.0 ± 1.6 SE and 45.6 ± 1.5 μmol C L^−1^, respectively). However, both depths showed similar dynamics, with relative maxima of DOC at noon and also at midnight at the surface (**Fig. 2A**) and a slightly higher CV in the fish layer (8.1% vs. 5.7%). The apparent hourly rates of DOC production (from 8 am to 12 pm at both depths) and consumption (from 12 am to 8 am at the surface and from 12 pm to 4 pm at 550 m) were similar within each layer: ca. 1.3 μmol C L^−1^ h^−1^ at the surface and 2.0 μmol C L^−1^ h^−1^ at the fish layer. Regarding the fluorescent DOM fraction, the protein (Tyrosine)-like component C4 was on average one order of magnitude higher at the surface than at 550 m, although it showed more variability at depth (**Table S1**).

**Fig. 1.**
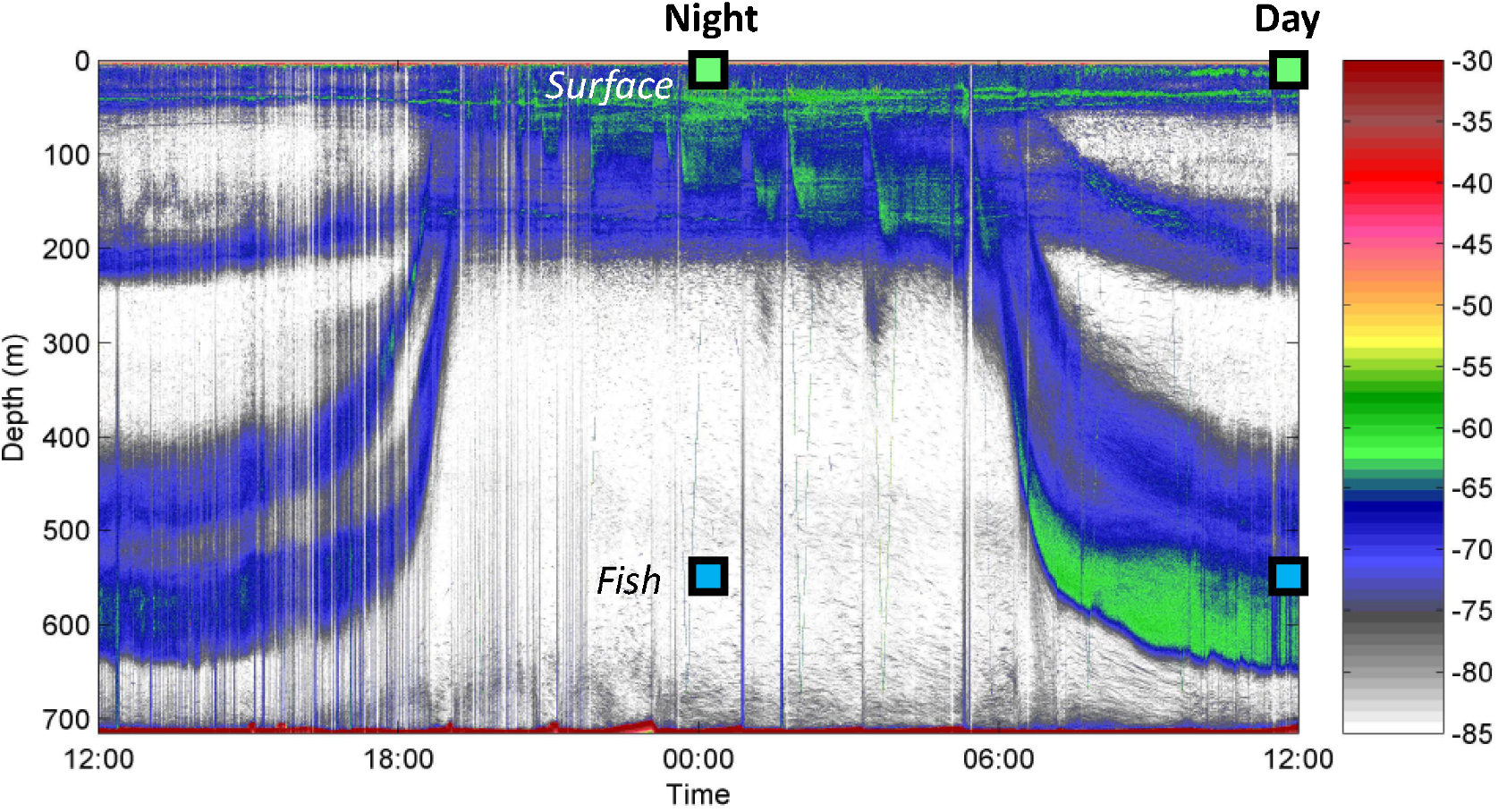
Echogram from March 5^th^ to 6^th^ 2016 at the study site showing 2 scattering layers of mesopelagic fish performing diel vertical migration: up to the surface at night and down to deep waters during daytime. Squares indicate the depth and time of water collection for the incubation experiments. Surface and mesopelagic (Fish) depths are represented in green and blue, respectively, for coherence with subsequent figures. Colour scale indicates backscattering strength (Sv, dB).

**Fig. 2.**
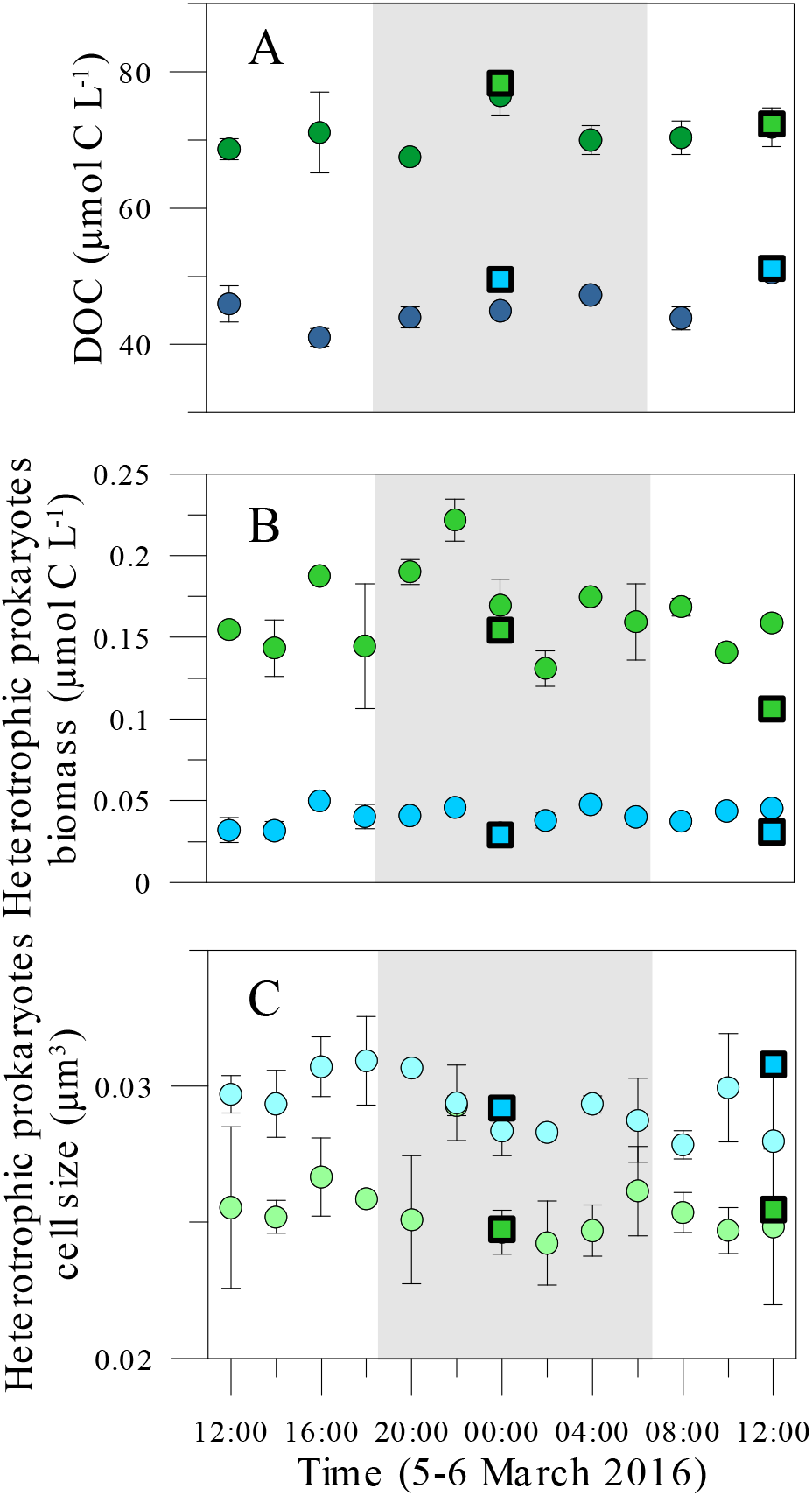
Variability of mean DOC concentration **(A)** and heterotrophic prokaryoplankton biomass **(B)** and cell size **(C)** in two layers of the study site: upper (0-25 m) and mesopelagic occupied by fish during daytime (450-600 m) during the 24 h sampling. Surface and mesopelagic (Fish) depths are represented in green and blue, respectively. Squares indicate initial values at the onset of the experimental incubations. The gray area represents nightime hours at the date of sampling. Error bars represent the standard error of the mean (average of 0 and 25 m in the upper layer and 450, 550 and 600 m in the mesopelagic one).

The abundance of HP at the surface (mean 4.31 ± 0.17 × 10^5^ cells mL^−1^) was also one order of magnitude higher than at 550 m (mean 9.43 ± 0.39 × 10^4^ cells mL^−1^), but varied similarly with no clear diel patterns. Although their size was 22% ± 8% larger in the mesopelagic (mean values of 0.034 and 0.028 μm^3^ at the fish and surface layers, respectively, **Fig. 2C**), the corresponding biomass was driven mostly by changes in abundance, averaging 1.91 ± 0.09 μg C L^−1^ at the surface and 0.50 ± 0.02 μg C L^−1^ at 550 m. **Fig. 2B** (in μmol C L^−1^ for comparison with **Fig. 2C**) shows that HP biomass was equally variable at both layers (CV 16.0 vs. 16.3%). In situ apparent or net growth rates based on changes in HP cell size were of 0.15 and 0.10 d^−1^ at 0 and 550 m, respectively. These estimated specific growth rates changed cyclically over the 24 h cycle, especially at the surface. Two maxima, at 20:00 and 8:00, were found at the surface layer while the maximum at 550 m was observed at 16:00 (**Fig. S2**).

### Experimental incubations of surface and deep samples

With initial concentrations similar to ambient values (**Fig. 2**), DOC was consumed in the first 2-3 days in the predator-free experiments (**Fig. 3**), albeit at different daily rates (**Table 1**), followed by net production after day 4 especially in the Surface incubations. Minimum and maximum consumption rates were 0.32 and 2.69 μmol C L^−1^ d^−1^, found in the Fish and Surface Night experiments, respectively (FN and SN). Values in the other two experiments carried out with noon samples were below 1 μmol C L^−1^ d^−1^ (0.47 and 0.95 μmol C L^−1^ d^−1^, respectively in in FS and SD). The initial fluorescence intensity values of the component C4 in the experiments was higher but reflected the values measured concurrently in the water column (paired t-test, p>0.05, n=4). C4 showed a very consistent consumption pattern regardless of the layer and treatment. Heterotrophic prokaryotes in the Surface incubations consumed in 4 days 40 and 50% of the initial values during Night and Day respectively, while bacteria inhabiting deep waters consumed almost all of it (95-100 %) within the same time frame regardless of the sampling time (**Fig. 4**). Hereinafter we consider changes in C4 as representative of labile DOM dynamics. Day and night C4 consumption patterns did not show any significant differences within the same layer but they displayed significantly higher consumption rates in the Fish layer (ANOVA, p=0.004, post-hoc Fisher LSD test, **Figure 4** and **Table S1**).

**Table 1.** Mean ± SE values of specific growth rates (μ), DOC consumption rates or prokaryotic carbon demand (PCD, see the text), prokaryotic heterotrophic production rates (PHP) and prokaryotic growth efficiency (PGE) in the surface and fish layer incubation experiments performed at noon (Day) and midnight (Night). Rates were calculated for each period of exponential growth, also indicated in days. The same period was used for DOC consumption and biomass production rates. Also indicated are the maximum heterotrophic prokaryotes biomass reached within the incubation and the corresponding ratio of maximum to initial biomass (Max:t0 biomass ratio).

| Layer | Time | Period (d) | $\mu$ ($d^{-1}$) | DOC consumption rate ($\mu\text{mol C L}^{-1} d^{-1}$) | PHP rate ($\mu\text{mol C L}^{-1} d^{-1}$) | PGE (%) | Maximum HP biomass ($\mu\text{g C L}^{-1}$) | Max:t0 HP biomass ratio |
| --- | --- | --- | --- | --- | --- | --- | --- | --- |
| Surface | Day | 0-1.75 | $0.34 \pm 0.07$ | $1.0 \pm 0.5$ | $0.040 \pm 0.004$ | 4.2 | $2.69 \pm 0.20$ | 2.30 |
| | Night<sup>a</sup> | 0-2.25 | $0.18 \pm 0.02$ | $2.7 \pm 1.1$ | $0.047 \pm 0.006$ | 1.8 | $2.85 \pm 0.16$ | 1.80 |
| Fish | Day<sup>a</sup> | 0-2.75 | $0.16 \pm 0.04$ | $0.5 \pm 0.3$ | $0.020 \pm 0.004$ | 4.2 | $0.94 \pm 0.13$ | 3.27 |
| | Night | 0-2.25 | $0.09 \pm 0.01$ | $0.3 \pm 0.8$ | $0.010 \pm 0.001$ | 3.1 | $0.68 \pm 0.02$ | 2.31 |

<sup>a</sup>Vertically migrating mesopelagic fish present

**Fig. 3.**
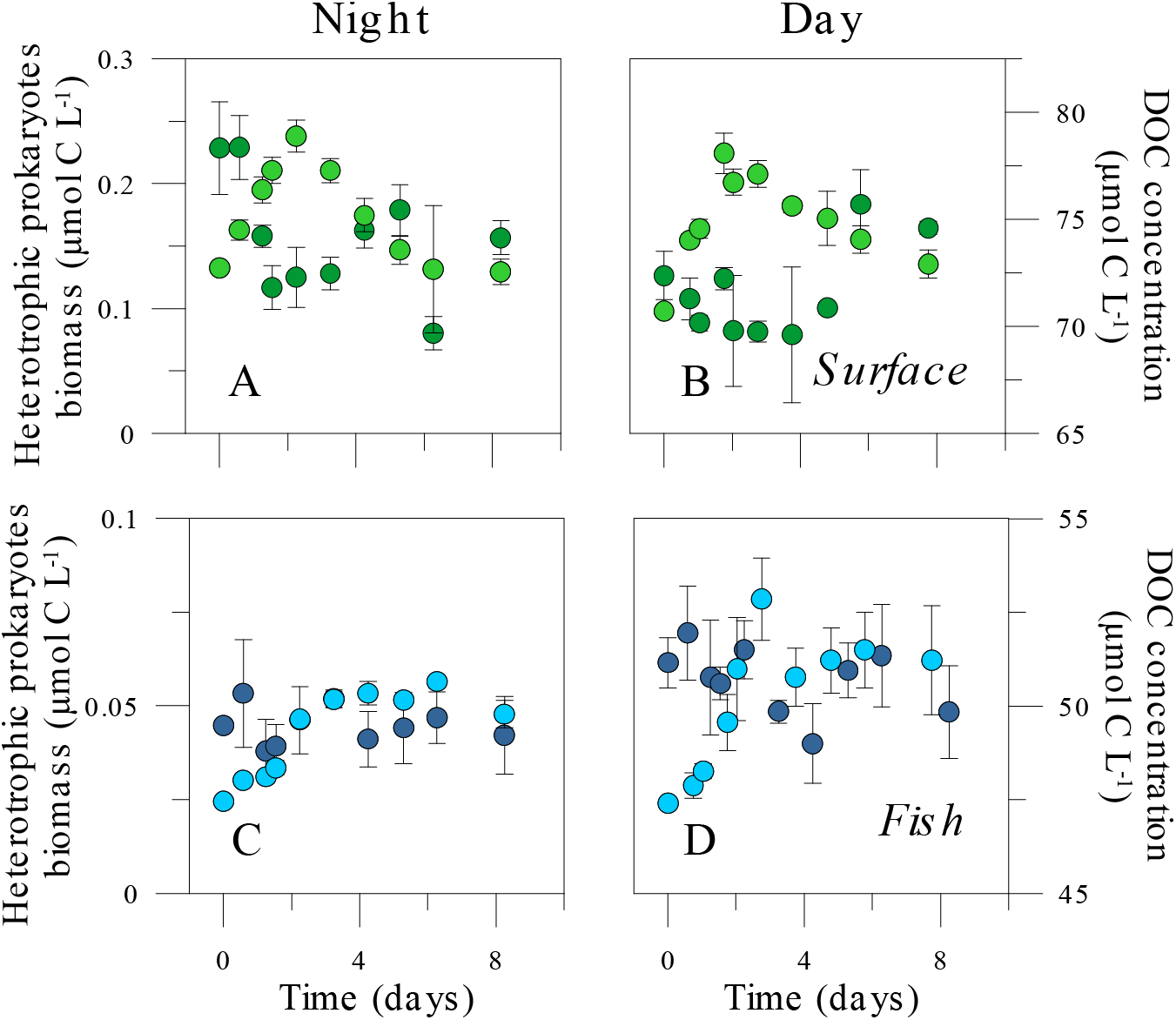
Dynamics of heterotrophic prokaryoplankton biomass (light colour) and DOC concentration (dark colour) in the predator-free experimental incubations of samples taken at noon (**A, C**) and at midnight (**B, D**) from the surface and the mesopelagic (Fish, 550 m depth) layers. Note the different scales for surface and mesopelagic water experiments. Error bars are standard errors of 3 replicates.

**Fig. 4.**
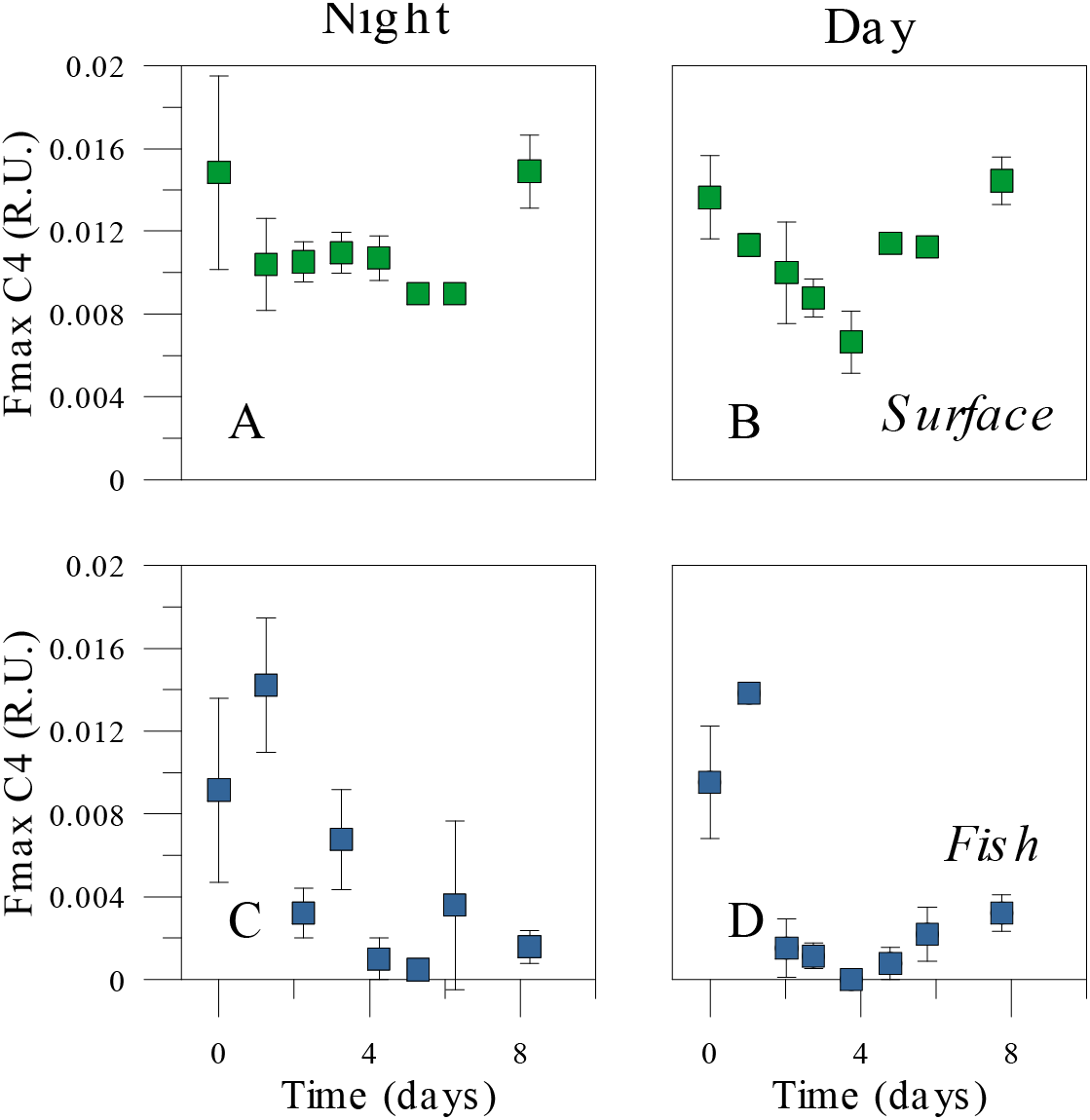
Dynamics of the concentration of the FDOM protein-like C4 component in the predator-free experimental incubations of samples taken at noon (**A, C**) and at midnight (**B, D**) from the surface and 550 m depth. Error bars are standard errors of 3 replicates.

Heterotrophic prokaryotes responses in the incubation experiments differed between depths and sampling times (**Fig. 3**). Consistent differences were found between the specific growth rates at both depths (**Table 1**), with μ being double in the Surface than in the Fish experiments. Within each layer, the Day μ values were also higher than the Night ones (t-tests, p=0.020 and p=0.060 at the Surface and Fish experiment, respectively, n=6). HNA cells always grew faster than their LNA counterparts resulting in increases in their relative contribution from 43-55% to 55-62%, more noticeable in the FD experiment. The mean size of the cells also increased substantially in the Fish experiments, from 0.027 to 0.060 μm^3^ in the FD incubation and from 0.028 to 0.047 μm^3^ in the FN one, while changes in cell size were much smaller in the Surface experiments, and virtually the same in both periods, from 0.026 to 0.037 μm^3^ (SD) and from 0.025 to 0.035 μm^3^ (SN) (**Fig. S2**). Consequently, cell size played an important role in the increase in biomass, especially in both Fish experiments (**Fig. 3C, D**). The biomass production rates of heterotrophic prokaryotes for the same periods of DOC consumption ranged 4-fold, from 0.010 to 0.047 μmol C L^−1^ d^−1^, mirroring the changes in the latter variable (**Table 1**). The rates of heterotrophic prokaryotes biomass production and DOC consumption were used for estimating prokaryotic growth efficiencies (PGE) in the four experimental incubations. PGE was uniformly below 5%, ranging from 1.8% (SN) to 4.2% (both SD and FD, **Table 1**). Following the pattern of in situ values, maximum HP biomass measured in the incubations was higher in the Surface than in the Fish experiments (**Fig. 3**, **Table 1**), although the increase ratios (i.e. the ratio of maximum to initial biomass, **Table 1**) were significantly higher in the Fish experiments with all data pooled (t-test, p=0.048, n=12).

## Discussion

### Diel interactions between DOM and heterotrophic prokaryotes

There is consensus that marine biota biomass and activity peak in the upper layers and decrease exponentially with depth, following the strong vertical gradients in physico-chemical properties ^31^. Heterotrophic prokaryotes inhabiting the Red Sea seem to challenge this view. Together with diel variations of standing stocks ^25^, this study), night- and day-initiated incubations of predator-free ambient assemblages with the DOM pool available at the time of sampling yielded surprising similarities in epipelagic and mesopelagic waters. A method for estimating division rates in cyanobacteria based in changes in cell size ^32, 33^ was adapted to obtain independent estimates of in situ net growth rates (**Fig. S2**). The diel variability in heterotrophic prokaryotes cell size showed the same pattern at the surface and the mesopelagic fish layer (**Fig. 2**), yielding low but comparable net growth estimates at the surface than at 550 m depth (0.10-0.15 d^−1^). However, it must be noted here that the already large prokaryotes inhabiting the mesopelagic fish layer were able to grow much bigger in the absence of protistan grazers (**Fig. S1**), confirming the results of a previous study conducted at the same site ^24^.

Diel cycles in biogeochemical properties and plankton biomass and activity in the upper ocean layers are reasonably well known, following diurnal changes in photosynthesis and food web processes ^4, 9^. No clear patterns were observed for in situ heterotrophic prokaryotes abundance or biomass, probably due to strong coupling between growth and mortality due to protistan grazing ^34, 35^ and/or viral lysis in the Red Sea (E. Sabbagh et al., in prep.), and in tropical waters in general ^36^. However, likewise early observations in the Caribbean ^37^ recently confirmed for this site ^25^, DOM concentrations displayed a coherent diel pattern suggesting different timing of production and consumption (**Fig. 1**). Confirming previous experiments ^38^, DOC consistently decreased in the first 2-3 days in both Surface and Fish samples, ranging from 0.7 to 6.1 μmol L^−1^, followed by net production after day 4 especially in the Surface incubations, coincident with a sharp decay in prokaryotic biomass (**Fig. 3A, B**). Indeed, the strong apparent uptake of surface DOC in situ during nighttime coincided with the highest DOC consumption experimental result, poiting to its labile nature, although the buildup of HP biomass was similar in both SD and SN experiments, resulting in lower specific growth rates and PGE values in the Night (**Table 1**). Labile DOC incorporation does not automatically inform us of its subsequent partitioning between metabolism (respiration) and growth (biomass production), as shown by Condon et al. ^39^ for DOM originated by jellyfish blooms. Since the alternation between light and dark periods within the incubator used for Surface seawater was common for SD and SN experiments, if photoheterotrophy ^40, 41^ afffected DOM dynamics, it should have been equally apparent in both Night and Day incubations, which were initiated with ony 12 h difference (**Fig. 1**). We therefore ruled out photoheterotrophic processes explaining the differences observed after 8 days of incubation. Rather, the quality of labile DOM at midday (including recent photosynthate) was probably higher than at midnight (perhaps with a greater contribution of DOM coming from sloppy feeding, ^42^, causing the significantly higher μ values (**Table 1**). That the quality of labile DOC at noon could have been higher was supported by a faster increase in bacterial cell size (**Fig. S1B**) and the contribution of HNA cells, which increased by 35% in SD compared with 15% in SN. The decrease in the protein-like C4 component was also more sustained in the SD experiment compared to SN after 4 days (13.2% vs. 5.3% was consumed daily during that period, **Fig. 4**) although the rates were virtually the same in the initial periods (2.75 and 2.25 days, respectively, **Table S1**). Changes in mesopelagic DOC concentrations and lability at diel (^25^, this study) and seasonal scales ^43^ in the central Red Sea support the recent claim that the diverse pool of DOM in the deep ocean fluctuates at timescales much shorter than previously thought ^44^. Since the conditions at the study site were hypoxic in most of the mesopelagic realm ^43^, the low in situ oxygen concentrations in the Fish water samples, consistent from 300 m downwards (0.69 ± 0.03 mg L^−1^) might have been supplemented by pre-filtration and sampling from the experimental bottles. Although we did not control for this potential artefact, the same protocol was followed for the FN and FD experiments. Therefore, the consistently higher values of DOM consumption, prokaryotic cell size, growth rates and efficiency in the Day compared with the Night incubation strongly support that the presence of fish indeed had a major impact on the microbial community.

Surface specific growth rate in the Day experiment was nevertheless notably higher than the 0.08 d^−1^ measured in a previous study carried out in November 2015 at the same location ^24^. The discrepancy cannot be explained by total DOC or chlorophyll *a* concentrations, but could instead be related to the availability of labile DOM compounds since C4 concentrations were 61% higher in March than in November (M. L. Calleja, pers. comm.). At a shallower, nearby site characterized by higher total and labile DOC concentrations, specific growth rates were still considerably higher, ranging from 0.79 to 1.75 d^−1 35^.

The daily rates of apparent DOC production and consumption based on changes in in situ concentration were virtually the same within each of the two layers compared, indicating no net accumulation. This is the expected result in oligotrophic regions at the short time scale of one day ^37, 45^. However, these rates were still ca. 50% higher in the mesopelagic zone resulting in apparent turnover of labile DOC of 23.6% d^−1^ in the mesopelagic compared with 14.7% d^−1^ at the surface, if we consider the measured diel variability and maximum concentrations in each layer. This finding conflicts with the contention that DOM is largely of refractory nature within the mesopelagic waters of the global ocean ^46^. The role of vertically migrating animals, zooplankton and fishes, as vectors of organic matter to deep layers complementary to the biological pump ^14^ has been recently recognized ^17, 18, 47^). In this regard, work at the study site has suggested DVM fishes as a transport mechanism supplying labile DOM that does not accumulate but fuels heterotrohic bacterial activity in mesopelagic waters^24^. Here we tested this hypothesis by further examining the fluorescence properties of DOM and its transformation by heterotrophic prokaryotes in experimental incubations with and without the fishes present. FDOM are useful tracers for biogeochemical processes in the dark ocean ^48, 49^. Fluorescence intensity of the two aminoacid-like fluorophores C3 and C4 decreased with depth (data not shown), indicating that these fluorophores were mainly produced autochthonously in surface waters. Both phytoplankton and bacteria are sources of tryptophan and tyrosine ^50^, while Urban-Rich et al. ^51^ have reported that grazing and excretion by zooplankton can also release material with amino acid-like fluorescence signals. Our results strongly suggest that DVM fishes can also provide C4 in the mesopelagic realm.

Contrary to the epipelagic zone, very few studies on the diel variability of DOM-heterotrophic prokaryotes interactions are available for deep waters ^25, 52^. Gasol et al. ^53^ suggested that mesopelagic prokaryotes in the subtropical NE Atlantic were as active as in the epipelagic. We demonstrate here not only that heterotrophic prokaryotes specific growth rates at 550 m were of the same order of magnitude than in surface waters, clearly challenging the most accepted view ^31, 54^, but that those rates were almost double at noon conditions, when the mesopelagic fishes were present, than at midnight, when the entire population was closer to the surface ^22^. From **Fig. 1** it is clear that the fishes were absent at midnight in the entire mesopelagic zone but their presence at 550 m had been established for ca. 4 h when the noon sampling took place. Specific growth rates were nevertheless lower than in the previous study (0.24 d^−1^, ^24^). Although seasonality of C4 in the deep scattering layer was less marked than at the surface, November 2015 was characterized by 82% higher C4 concentrations than in March 2016 (M. Calleja, pers. comm.). Altogether, these results point out to a major role of protein-like substances in determining the specific growth rates of heterotrophic prokaryotes throughout the water column, as recently found for nearby shallow waters in a seasonal study^35^. C4 fluctuated widely in the 24 h monitoring at 550 m depth (**Table S1**) and was also actively consumed in all our incubations, thus revealing a clearly labile nature. C4 was consumed faster in the Fish experiments, at 42.1% and 25.8% d^−1^ in FD and FN, respectively (**Fig. 4**, **Table S1**), than in the Surface ones (ca. 13% d^−1^ in both SD and SN). A similar relative consumption of protein-like FDOM (12% d^−1^), mostly occurring during the first 5 days, was measured by ^55^ in experiments conducted with marine surface waters. Our explanation is that fishes released DOM directly or it leaked from particles associated to the fish presence (e.g. fecal pellets). That DOM could have been delivered by sinking particles ^56^ not related to vertical migration would not explain the difference between the FD and FN experiments.

Cell size has been used as an indicator of the activity of heterotrophic prokaryotes ^57^. While in the Surface experiment growing HP cells were only slightly larger than at time 0 (11% larger size for both the SD and SN experiments), the cell size increase in the Fish experiment was dramatic, especially in the Day incubations (118% vs. 68% larger, **Fig. S2**). The contribution of bigger cells to the observed increase in HP biomass is not at all minor: had we used, as in many studies, a fixed cellular carbon content of 4 fg C cell^−1^ (corresponding to the initial mean cell size of 0.027 μm^3^ for the two depths and periods, **Fig. 2C**), maximum HP biomass in **Table 1** would have become 2.10 (SD), 2.32, (SN), 0.43 (FD) and 0.39 (FN) μg C L^−1^, i.e. between 42 and 54% lower than the actual values for the mesopelagic prokaryotes. We conclude that the presence of fishes in the mesopelagic zone during daytime resulted in significantly higher growth rates of markedly larger cells. As a consequence, changes in abundance were exacerbated when considering biomass units. The maximum biomass of heterotrophic prokaryotes that could be sustained by extant DOM concentrations was significantly higher than the initial value in both Fish experiments (**Table 1**). Altogether, these results point out to substantial inputs of labile DOM during daytime at the mesopelagic fish layer that are rapidly mobilized by large bacterial taxa. The archaeon *Nitrosopulimus maritimus*, which makes up much of the heterotrophic prokaryoplankton biomass at these depths ^58^, were apparently not the main responders to these DOM hotspots, since their contribution to total numbers at the end of a similar incubation dropped from 50% to 3% ^24^. The typical size of these Thaumarchaeota is small ^59^, so it is unlikely that they were the dominant groups growing in our Fish incubations after 2 days (**Fig. S2C**, **D**). Besides excreting ammonia that boosts its oxidation by Thaumarchaeota (formely Chrenarchaeota) ^60^, mesopelagic fish thus seem capable to fuel the metabolism of large, copiotrophic bacteria.

Prokaryotic growth efficiencies are typically low in open ocean, oligotrophic environments ^61, 62^. Most of these measurements refer to the epipelagic zone, where photosynthesis takes place. Here we provide more information from deeper layers, where export processes are more important. The recently reported low PGE values at this Red Sea site (1.6-3.4%, ^24^ are confirmed by this new study, while higher values (2.5-12.8%) were recorded in a shallow, richer bay located a few km south ^35^. Few studies have estimated the vertical variability in PGE values, but those that have usually depict lower values with depth ^62, 63^, related to the increased presence of refractory DOM compounds ^46^ or to the higher dilution of the labile ones ^64^. Notably, the estimated growth efficiency of heterotrophic prokaryotes in our Day experiments was exactly the same in Surface and Fish water (4.2%), which can only be explained by the existence of labile DOC of similar quality within both layers, resulting from photosynthesis at the surface and fish-mediated export in the mesopelagic layer. PGE in the mesopelagic fish layer was significantly higher than at shallower depths in the experiment conducted at noon in November 2015 ^24^. However, when averaging our new two estimates, the mean PGE value at 550 m (3.6%) was still 22% higher than at the surface, strongly supporting the presence of high quality DOM hotspots ^24^ in the deep scattering layer.

Although this study is limited to a single 24 hour period, it builds on the previously demonstrated relatively high growth of mesopelagic heterotrophic prokaryotes ^24^ by comparing the outcome of deep samples incubations taken only 12 hours apart, at midnight and noon. We confirm that the Red Sea mesopelagic zone is not a permanently impoverished environment but subject to daily inputs of labile DOM compounds similarly to the epipelagic layers. This novel process driven by mesopelagic fishes, which complements other recently discovered sources of deep organic carbon ^14, 65, 66, 67^ seems to have been overlooked due to the tight coupling between the components of microbial food webs ^68^. If vertically migrating fishes are able to fuel an active and distinct community (T. Huete-Stauffer et al., submitted) of heterotrophic prokaryotes in the mesopelagic layer of the Red Sea, we might expect this fast DOM flux to be widespread. The mesopelagic Red Sea has an unusually high temperature, therefore the effect of colder conditions on fish DOM-microbial interactions remain to be explored. The implications for global biogeochemical cycling would also vary depending on the actual biomass of mesopelagic fishes and the fraction performing DVM ^20^, yet its impact may increase as deep waters warm up ^69^. That these small fishes seem able to sustain the microbial communities inhabiting the twilight zone also may help reconcile current discrepancies between carbon pools and fluxes in the global ocean.

## METHODS

### Environmental sampling

We occupied one station located 13.4 km offshore to the north of King Abdullah Economic City, Saudi Arabia (lat 22.46°N, lon 39.02°E) between midday March 6^th^ and midday March 7^th^, 2016 ^24, 25^. In situ monitoring and sampling was conducted on board of RV Thuwal. Continuous acoustic measurements in order to locate the position of the vertically migrating mesopelagic fishes were recorded with a Simrad EK60 38 kHz echosounder mounted on the ship’s hull. From noon on March 6^th^ until the same time on the following day we conducted CTD casts every 2 hours. At each cast we sampled discrete depths in the water column with Niskin bottles mounted on a Rosette sampler, ranging from the surface to 650 m depth. Water filtered through pre-combusted Whatman GF/F filters was collected for analyzing DOC bulk concentrations and fluorescent DOM (FDOM) properties (40 mL pre-combusted glass vials). Unfiltered water was collected for characterizing the community of heterotrophic prokaryotes (2 mL cryovials).

Hourly in situ apparent DOC production and consumption rates were estimated as the largest difference between DOC concentration in consecutive sampling times that showed a consistently increasing and decreasing trend, respectively. The same approach was used for estimating the apparent biomass production of heterotrophic prokaryotes over the diel cycle.

### Experimental incubations

10 L of seawater from the surface and 550 m depth were collected in the midnight and noon casts on March 7^th^ for conducting the experimental incubations of DOC consumption, change in FDOM and heterotrophic prokaryotes biomass response. In order of remove protistan grazers and planktonic organisms larger than bacteria and archaea, water was gently filtered through pre-combusted Whatman GF/C filters (142 mm, nominal pore size 1.2 μm) and used to fill 3 × 2 L acid-cleaned polycarbonate bottles, which were subsequently incubated at the in situ temperature and light regimes (darkness for 550 m samples), so that the possible role of photoheterotrophs in the processing of DOM was included. Removal of heterotrophic prokaryotic cells by filtration was minor (83% ± 7% SE of the initial abundance was retrieved in the water used for the incubations) and mean cell size was virtually unaffected (2.6% ± 1.0% smaller biovolume than inthe unfiltered water). However, filtration eliminated most *Synechococcus* and *Prochlorococcus* cyanobacteria as well as virtually all the larger protistan grazers of heterotrophic prokaryotes, since the mean abundance of heterotrophic nanoflagellates in the GF/C filtrate was 1.5% (E. I. Sabbagh, pers. comm.). We are therefore confident that no extra sources of DOM were included in our experiments. Subsamples were taken twice per day on the first 2 days, then daily until day 6 and finally at day 8. DOC and FDOM subsamples from the incubations were gravity-filtered through pre-rinsed 0.2 Millipore polycarbonate filters. We will occasionally use the codes S and F to refer to the incubations made with water from the Surface (0 m) and the Fish layer (550 m), respectively, followed by D or N to refer to the period of sampling (Day or Night): SD, SN, FD, FN.

### DOC analysis

Samples for DOC were acidified with H_3_PO_4_ and kept in the dark at 4 °C until analysis by high temperature catalytic oxidation at the laboratory. All glass material used was acid cleaned and burned (450°C, 4.5 h). Consensus reference material of deep sea carbon (42–45 μmol C L^−1^ and 31-33 μmol N L^−1^) and low carbon water (1-2 μmol C L^−1^), provided by D. A. Hansell and W. Chen (Univ. of Miami) was used to monitor the accuracy of our DOC concentration measurements. The analytical error of DOC concentration was 1.4 μmol L^−1^.

### DOM fluorescence measurements and PARAFAC modeling

FDOM samples were stored at 4°C before being analyzed (within 2 days after the completion of the cruise and incubation sample collection). UV-VIS fluorescence spectroscopy was measured using a HORIBA Jobin Yvon AquaLog spectrofluorometer with a 1 cm path length quartz cuvette. Three dimensional fluorescence excitation emission matrices (EEMs) were recorded by scanning with an excitation wavelength range of 240-600 nm and emission of 250-600 nm, both at 3 nm increments and integrating at 8 seconds. To correct and calibrate the fluorescence spectra post-processing steps were followed according to Murphy et al. ^70^. Briefly, fluorescence spectra were Raman area (RA) normalized by subtracting daily blanks that were performed using Ultra-Pure Milli-Q sealed water (Certified Reference, Starna Cells). Inner-filter correction (IFC) was also applied according to McKnight et al. (2001) RA normalization, blank subtraction, IFC and generation of EEMs were performed using MATLAB (version R2015b).

A total of 165 samples for DOM fluorescence were collected (81 from 7 vertical profiles and 84 from the experimental incubations). The EEMs obtained were subjected to PARAFAC modeling using DOMFluor Toolbox ^71^. Before the analysis, Rayleigh scatter bands were trimmed. A four-component model was validated using split-half validation and random initialization ^71^: peak C1 at Ex/Em 240(325)/ 407 nm, peak C2 at Ex/Em 258(390)/492 nm, peak C3 at Ex/Em 240/337 and peak C4 at Ex/Em 276/312 nm. C1 corresponds to peak M ^72^ and is comparable to component 2 identified by Català et al. ^49^. C2 represents a combination of peaks A and C ^72^ and is comparable to component 1 in ^49^. C3 corresponds to peak T ^72^, attributed to tryptophane, and is comparable to component 3 in ^49^. C4 corresponds to peak B ^72^, attributed to tyrosine, and is comparable to component 4 in ^49^. The maximum fluorescence (Fmax) is reported in Raman units (RU).

### Heterotrophic prokaryotes abundance and biomass

Triplicate samples (1.8 mL) for estimating the abundance of heterotrophic bacteria and archaea in situ and in the experimental incubations were fixed with 1% paraformaldehyde and 0.05% glutaraldehyde, deep frozen in liquid nitrogen and stored at −80°C until analysis. Once thawed, 400 μL aliquots were stained with SYBR-Green run in a BD FACSCanto II flow cytometer for estimating the abundance of low (LNA) and high (HNA) nucleic acid content cells as detailed in Gasol and Morán ^73^. The few cyanobacteria present were easily distinguished in the surface samples due to their autofluorescence red signal because of the presence of chlorophyll *a*. Absolute abundances were estimated based on time and the actual flow rates, which were calibrated daily using the gravimetric method. The right angle light scatter or side scatter (SSC) signal relative to the value of 1 μm fluorescent latex beads added to each sample was used to estimate the cell diameter according to Calvo-Díaz and Morán ^74^. LNA and HNA cell numbers were summed to estimate the total abundance and their specific cell sizes averaged to obtain the mean cell size of the heterotrophic prokaryote community at both depths and different times. Assuming spherical shape, the mean cell size (biovolume in μm^3^) was converted into cellular carbon content following Gundersen et al. ^75^. Heterotrophic prokaryotes biomass was then calculated as the product of cell abundance and mean cellular carbon content.

### Growth rate estimates

In situ apparent or net growth rates of the heterotrophic prokaryote assemblage at the surface and the mesopelagic fish layer were estimated from changes in biomass (μg C L^−1^) resulting from changes in abundance and mean cell size over 24 h. Net growth rates (μ, in units h^−1^) were calculated as:

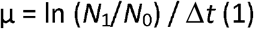

where *N*_1_ is the final biomass, *N*_0_ is the initial biomass and Δ*t* is the time interval (2 h). We modeled the overall daily growth rate from Eq. (1) using the size distribution of the organisms with the R package ssPopModel, which included a modified version of the size-structured matrix population model originally developed by Sosik et al. ^32^. Matrix population model assumption is that changes in size distribution are only related to growth and division of the cells. We adapted and simplified the application of this function as described by Hunter-Cevera et al. ^33^ for *Synechococcus* cyanobacteria to be used with heterotrophic prokaryote cells.

Specific growth rates in the incubations were calculated as the slope of the ln-transformed total abundance vs. time for the linear response period, equivalent to the phase of exponential growth (usually lasting between 2 and 3 days).

### Prokaryotic growth efficiency

Prokaryote heterotrophic production (PHP) in the midnight and midday incubations was estimated as the rate of increase in bacterial biomass during the exponential phase of growth. Prokaryotic carbon demand (PCD, i.e. the sum of heterotrophic prokaryotes production and respiration) was approached by the consumption rate of DOC during the same period. Prokaryotic growth efficiency (PGE) was therefore calculated as the ratio of PHP to PCD.

### Statistical analyses

Model I or ordinary least squares (OLS) linear regressions for estimating specific growth rates were done separately for each replicate, using a common period for each experiment. Differences between treatments and/or depths were assessed with one way ANOVAs and Fisher least significance (LSD) post-hoc tests. General relationships between variables were represented by Pearson’s correlation coefficients. Statistical analyses were done with JMP and STATISTICA software packages.

## ACKNOWLEDGMENTS

We are greatly indebted to the crew of RV Thuwal and the rest of the personnel from the Coastal and Marine Resources (CMOR) Core Lab at KAUST for their assistance during field work. Besides participating in the sample collection M. Viegas helped us with the rest of the work in the Red Sea Research Center (RSRC) lab. We are also grateful to past and current members of the Microbial Oceanography and Biogeochemistry lab at the RSRC.

X.A.G.M. conceived the research, led the experiment design, data analysis and wrote the paper. F.C.G. modeled in situ growth rates. A.R. performed the acoustic research. L.S. and N. A-O. analyzed the heterotrophic prokaryotes. X.I. contributed to the interpretation of results. M.L.C. was responsible for DOC and FDOM measurements, contributed to experimental design and data analysis. F.C.G., A.R., X.I. and M.L.C. also contributed to writing.

## Competing interests

The authors declare that they have no financial or non-financial competing interests.

## Notes

### Competing Interest Statement

The authors have declared no competing interest.

### Summary of Updates

Text and figure captions updated for clarity. More methodological details provided.

