## Supplemental Figures 1 and 2 and Table 1 for "Diel dynamics of dissolved organic matter and heterotrophic prokaryotes reveal enhanced growth at the ocean’s mesopelagic fish layer during daytime"

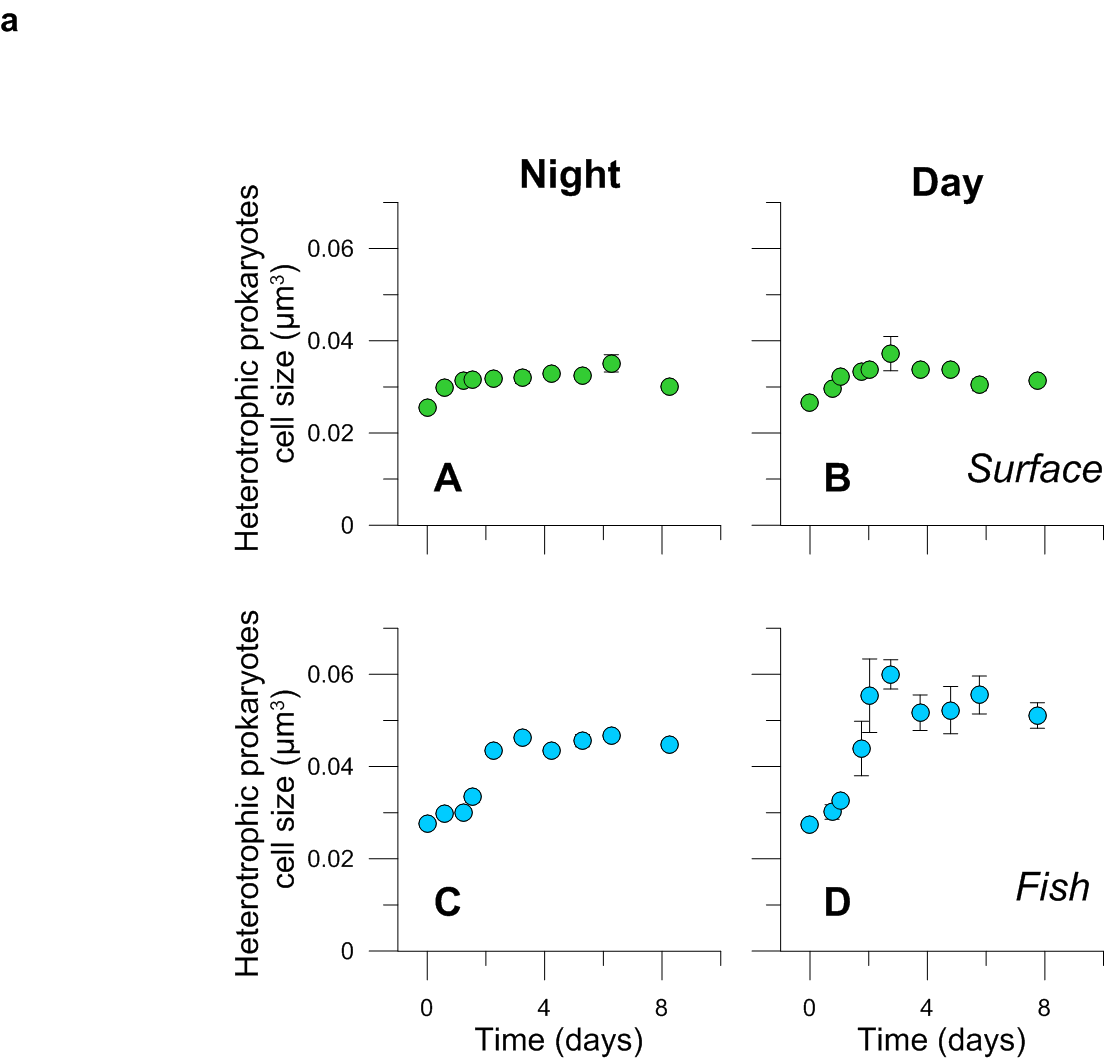

**Fig. S1**. Dynamics of heterotrophic prokaryoplankton cell size in the predator-free experimental incubations of samples taken during daytime (A, C) and nighttime (B, D) from the surface and 550 m depth at KAEC stations.

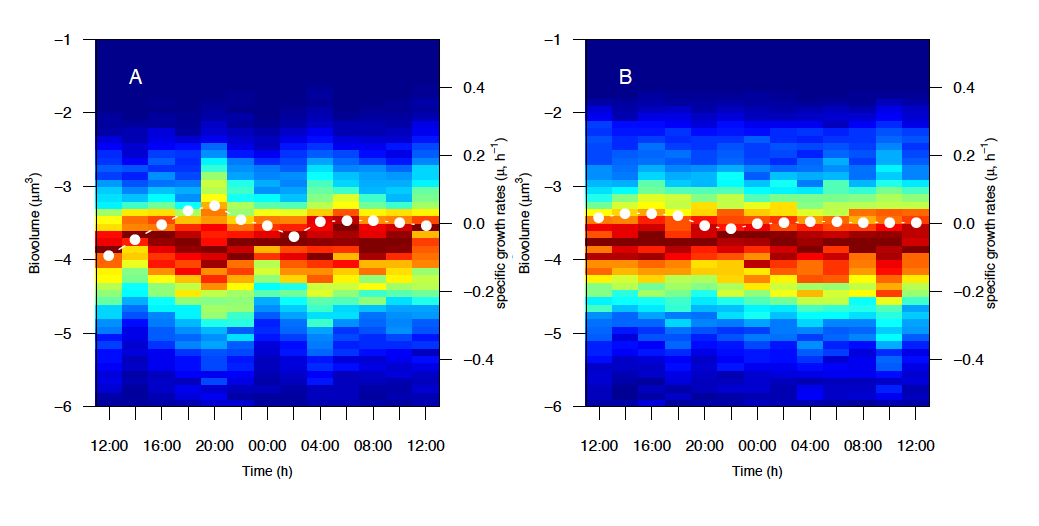

**Fig. S2**. Size distribution (µm^3^) of heterotrophic prokaryotes along the 24 h sampling at the surface (A) and 550 m depth (B). The color code represents the normalized fraction of cells (0, blue; 1, red) that falls on each size class. The white line represents the specific growth rates (µ, h^-1^) based on changes in heterotrophic prokaryotes biomass with time.

**Table S1.** Mean ± SE in situ concentrations of C4 and consumption rates in the experiments performed with noon (Day) and midnight (Night) samples (see **Fig. 3**). The period of calculation coincided with the exponential growth phase of heterotrophic prokaryotes.

| **Layer** | **C4 concentration**  **( x 10^-3^ R.U.)** | **Time** | **Period**  **(d)** | **C4 consumption rate**  **(x 10^-3^ R.U. d^-1^)** |
| --- | --- | --- | --- | --- |
| Surface | 12.26 ± 0.92 | Day | 0-2.75 | 1.73 |
|  |  | Night | 0-2.25 | 1.99 |
| Fish | 1.82 ± 1.04 | Day | 0-2.75 | 4.01 |
|  |  | Night | 0-2.25 | 2.36 |
